# Functional traits of carabid beetles reveal seasonal variation in community assembly in annual crops

**DOI:** 10.1101/2021.02.04.429696

**Authors:** Ronan Marrec, Nicolas Gross, Isabelle Badenhausser, Aurélie Dupeyron, Gaël Caro, Vincent Bretagnolle, Marilyn Roncoroni, Bertrand Gauffre

## Abstract

1. Trait-based community assembly studies have mostly been addressed along spatial gradients, and do not consider explicitly a fundamental dimension governing community assembly, the time. Nevertheless, such consideration seems particularly necessary in systems in which organisms have to face regular disturbances and rapid changes in vegetation phenology, such as in intensively managed farmlands.
2. In this study, we aimed at understanding how the functional diversity of carabid beetle communities varied across the growing season in response to crop type. We tested three alternative hypotheses on mechanisms underlying the community assembly.
3. We used data from a long-term monitoring conducted over nine years in an intensively-managed farmland in central western France, in a total of 625 fields. First, we measured morphological traits related to body size, dispersal mode, and resource acquisition on the 13 dominant carabid species (> 85 % of all trapped individuals) and identified three independent dimensions of functional specialization within our species pool along axes of a PCA and highlighted key traits for community assembly. Second, we evaluated the community assembly temporal dynamics and the impact of habitat filtering and niche differentiation in the different crop types with time, using linear mixed-effects models.
4. We showed that functional species assembly of carabid beetle communities occurring in crop fields varies importantly intra-annually, with strong variations in these dynamics depending on crop type and crop phenology. Each crop acted as a filter on carabid communities for body size and resource-acquisition traits, and functional differentiation between crops increased with time. We did not find any evidence of habitat filtering on traits related to dispersal mode.
5. Our results emphasize the major role of crop phenology but also disturbances involved by agricultural practices such as crop harvesting on changes in community assembly, likely due to seasonal and inter-annual redistributions of species in agricultural landscapes in response to such changes. The temporal dimension cannot be ignored to understand the assembly of local carabid communities in farmlands.

## 1. INTRODUCTION

Trait-based approaches are considered as one of the most prominent tool for the study of community assembly for both plants (e.g., Kraft et al., 2008; Le BagousseūPinguet et al., 2017) and animals (e.g., Gaüzère et al., 2015; Le Provost et al., 2017). Deterministic processes that shape plant and animal communities can be broadly separated into two distinct families with opposite effects on species assemblage. First, habitat filtering corresponds to any process that selects species with similar trait values (Keddy, 1992; Maire et al., 2012). At the community level, habitat filtering leads to trait convergence toward an optimal trait value that matches the local abiotic/biotic environment (Grime, 2006). Second, niche differentiation (e.g., limiting similarity, MacArthur & Levins, 1967) favours individual species with contrasted traits values (Maire et al., 2012). At the community level, niche differentiation can lead to high trait diversity by promoting species exploiting locally contrasted resources (HilleRisLambers et al., 2012). Trait-based community assembly studies have mostly been addressed along spatial gradients (e.g., Le Bagousse□Pinguet et al., 2017; Vanneste et al., 2019). While these studies are useful to detect how environmental conditions shape the functional structure of communities, they do not consider explicitly the temporal dynamics of communities and their environments.

Ecological communities face recurrent disturbances which may create transient community dynamics (Mouquet, et al., 2003) and instable equilibrium states (Scheffer et al., 2001). This source of variation may blur our ability to detect how trait differences between species determine community assembly. For instance, the trait diversity within communities has been shown to increase with time after disturbance (Fukami et al., 2005). In addition, most organisms are characterized by seasonal dynamics which may have profound implications for the study of community assembly (Fitzgerald et al., 2017; Habel et al., 2018). How disturbance interacts with seasonal dynamics of organisms in real situation is however largely unknown although assembly time and disturbance regime are theoretically predicted to interact and determine the relative importance of stochastic *vs.* deterministic processes on community assembly (Mouquet et al., 2003).

In agricultural landscapes, wild organisms have to face regular disturbances, such as direct destruction of their habitat, regular ploughing, and chemical treatment application, which strongly alter their abundance and taxonomical and functional diversities (Newbold et al., 2015). This is typically the case of carabid beetle communities which represent a functionally diverse guild of predators (Kromp, 1999). Carabid beetles exhibit a large interspecific variation in body size and in habitat and feeding preferences (Kotze et al., 2011; Lövei & Sunderland, 1996; Ribera et al., 2001). However, land-use intensification tends to reduce functional diversity of carabid communities (Woodcock et al., 2014), and for instance, select for smaller carabid species with higher dispersal abilities (Ribera et al., 2001) and lower feeding niche breath (Winqvist et al., 2014). By selecting species with similar traits, we could predict that the dynamic of species assembly within carabid communities is random due to high functional equivalence between species (Chesson, 2000; Gross et al., 2015; Hubbell, 2005). However, carabid beetle community structure has been shown to vary among different crop types (Eyre et al., 2013; Marrec et al., 2015). For instance, grassland habitat may offer stable habitat over time within agricultural landscape, and has been shown to promote functional diversity for plants and arthropods (Le Provost, Badenhausser, Le Bagousse-Pinguet, et al., 2020; Le Provost, Badenhausser, Violle, et al., 2020; Le Provost et al., 2017). In addition, carabid beetles may be sufficiently mobile (Ribera et al., 2001) to develop temporal strategy of habitat use, especially in response to seasonal environmental changes such as crop rotations (Holland et al., 2009; Marrec et al., 2015; Thomas et al., 2001). However, how such strategies and environmental influences affect carabid functional assembly remains unknown. Understanding how carabid communities change over time within and between crop types may help to design landscape-level management practices aiming at supporting key ecosystem services such as pest control, essential for global food production (Woodcock et al., 2014).

Here, we tested how the functional diversity of carabid beetle communities varied across the growing season in response to crop type. We used data from a long-term monitoring design conducted over nine years in an intensively-managed farmland (covering ca. 430 km^2^ in central western France). Carabid communities have been surveyed in a total of 625 fields from 2005 to 2013 over the spring-summer growing season. We first evaluated how morphological traits co-vary between species in order to identify independent dimensions of functional specialization within our species pool and highlight key traits for community assembly (Maire et al., 2012). We then tested three alternative, but non-exclusive, hypotheses on mechanisms underlying the community assembly of carabid communities: *Hypothesis 1,* community assembly is driven by random processes due to high functional equivalence between carabid species (Hubbell, 2005). In that case changes in community diversity are mostly due to the seasonal phenology of carabid communities and apparent random redistribution of individuals across communities. That would result in no differences in community structure between different crop types at a given time.

*Hypothesis 2,* farmland carabid beetles are adapted to high disturbance rate (Marrec et al., 2015) and characterized by fast assembly time (Mouquet et al., 2003). In that case, they are able to follow high temporal fluctuations of crop distribution, phenology, and resources. That would result in the fact that each crop may act as a filter on carabid communities and that functional differentiation between crops increases with time during the crop growing season (Fukami et al., 2005).

*Hypothesis 3,* functional diversity of carabid communities is higher in grasslands than in annual crops because grasslands show higher stability over time and present a more complex and diverse vegetation (Pakeman & Stockan, 2014). That would result in high functional diversity for all trait dimensions in grasslands during the entire season.

## 2. MATERIALS AND METHODS

### 2.1. Study area

The study was conducted in the Long Term Ecological Research “Zone Atelier Plaine & Val de Sèvre” area (LTER ZA-PVS) located in central western France (46°23’N, 0°41’ W). It is a farmland area of ca. 430 km^2^ mostly dedicated to cereal crop production. Since 1994, land use has been recorded annually for each field (~ 13 000 fields) and mapped with a Geographical Information System (ArcGis 9.2 - ESRI Redlands, CA, USA). From 2005 to 2013, land cover was dominated by annual crops, mostly winter cereals (36.9 % ± 0.4 of the total area), oilseed rape (10.1 % ± 0.7), and sunflower (10.8 % ± 0.5). Other crop types accounted for 18.2 % ± 3.4 of the land use. Temporary (sown with pure grasses or with mixed grasses with or without legume species and < 6 yr-old) and permanent grasslands (> 5 yr-old) represented 8.5 % ± 0.4 of the total area, and artificial grasslands (sown with pure legume species and < 6 yr-old; exclusively alfalfa in the study site) represented 3.4 % ± 0.3. Other main land uses were urban areas (9.3 % ± 0.3) and woodland (2.9 % ± 0.1) (Bretagnolle et al., 2018a).

### 2.2. Carabid beetle sampling

From 2005 to 2013, carabid beetles were sampled in the five dominant crop types in the study region (i.e., alfalfa, grassland, oilseed rape, sunflower, and winter cereals). The surveyed fields were randomly selected within the study area (see Appendix S1 for a full description of the data set). The comparative crop calendar of these crops in the study area can be found in Fig. 1. We used pitfall traps, the standard method to estimate carabid beetle abundance-activity (AA) during their activity period (Thiele, 1977). One to seven trapping sessions were conducted per field in a given year.

**Fig. 1.**
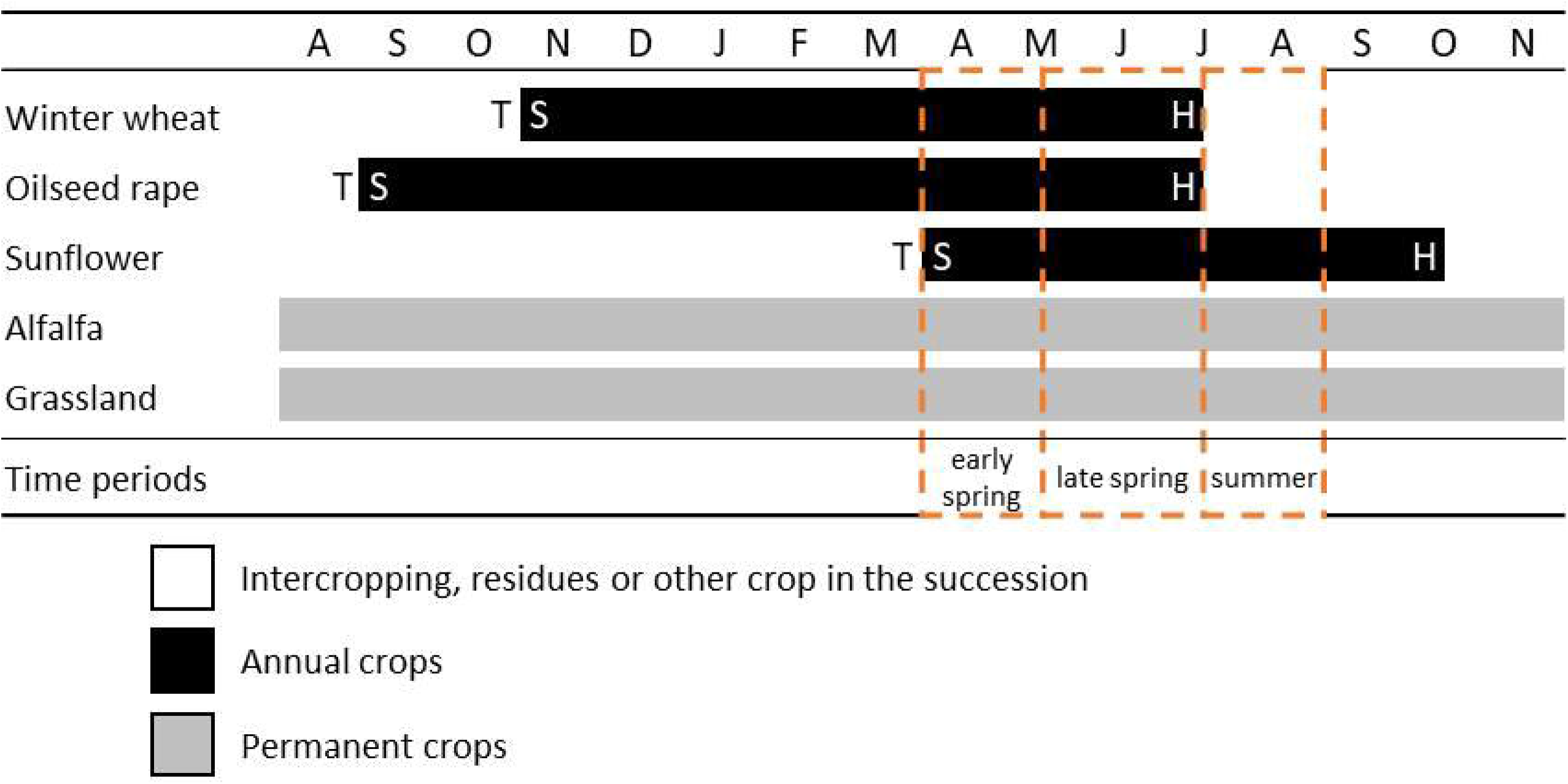
Crop calendar in the study area of the five dominant and sampled crop types. The dashed, orange boxes delineate the three time periods of sampling: early spring, April 1 to May 15; late spring, May 15 to July 10; summer, July 10 to August 30. T: soil tillage; S: sowing; H: harvesting.

Three pitfall traps were placed between 10 and 30 m from the field margin and at 10 m from each other. Traps were filled with a 50 % preservative solution of ethylene glycol (2005 to 2010), monopropylene glycol (2009 and 2010), or ethanol (2011 to 2013) (Bretagnolle et al., 2018b). The different preservative solutions used may affect catch probability (Skvarla et al., 2014) but the differences in AA among crops are robust for this bias (Marrec et al., 2015). Pitfall traps were left in place for five (2005-2010) or four (2011-2013) trapping-effective days and, for a given year, were set up at the same location for all sessions (see Appendix S1 for a complete description of the dataset). Carabid beetles were stored in the lab in a 96° ethanol solution and identified at the species level following Jeannel (1941, 1942). Species names follow *Fauna Europaea* (de Jong et al., 2014). Data from all the pitfall traps were aggregated per field and date, and used as the statistical unit in the following analyses. Overall 1,209 carabid communities were obtained from 625 fields and five crop types.

### 2.3. Species selection and trait measurements

In this study, we considered the 13 dominant carabid beetle species which accounted for 87.8 % of the catches in pitfall traps along the nine trapping years (57,409 individuals in total). The same 13 species were among the most abundant species in each year of the study: *Poecilus cupreus* (32.8 %), *Brachinus sclopeta* (19.0 %), *Anchomenus dorsalis* (13.6 %), *Calathus fuscipes* (4.3 %), *Nebria salina* (4.2 %), *Brachinus crepitans* (4.1 %), *Pterostichus melanarius* (2.4 %), *Harpalus dimidiatus* (2.2 %), *Harpalus distenguentus* (1.5 %), *Amara consularis* (1.4 %), *Pseudoophonus rufipes* (1.3 %), *Microlestes minutulus* (0.6 %), and *Microlestes maurus* (0.4 %). Morphological traits were measured on these 13 selected dominant species according to standardized protocols (Le Provost, Badenhausser, Le Bagousse-Pinguet, et al., 2020). Twelve individuals per species and per sex from our local species collection were measured. Measured individuals were selected randomly from 2011 and 2012 samples irrespectively of the crop type from which they have been trapped.

On each individual, we measured three sets of traits related to body size, movement ability, and resource acquisition that describe leading dimension of forms and functions in arthropods (Le Provost, Badenhausser, Le Bagousse-Pinguet, et al., 2020; Moretti et al., 2017). Body size is an important trait related to metabolic rate (Brown et al., 2004) and thermoregulation (Uvarov, 1977). For carabid beetles, body size is also a critical trait related to predation and pest control (Rusch et al., 2015). Movement ability traits may much vary between carabid beetles especially regarding their flight and running ability (Evans & Forsythe, 1984; Lövei & Sunderland, 1996). Finally, resource acquisition traits may also vary since carabid beetles have large range of feeding preferences ranging from granivory and herbivory to specialized carnivory (e.g., ectoparasitoids like many *Brachinus* species) (Lövei & Sunderland, 1996). The measured morphological traits were:

i. *Body size and shape-related traits:* body surface (*Bs*; mm^2^), measured as the sum of head, pronotum, and elytra areas; body length (sum of head, pronotum, and elytra lengths) *vs.* body width (abdominal maximum width) ratio (*Bl:Bw;* mm.mm^-1^); head length *vs.* head width ratio (*Hl:Hw;* mm.mm^-1^);
ii. *Mobility-related traits:* membranous wing surface (*Wg;* mm^2^); posterior leg length (*Lg;* mm); femur volume of the posterior leg (*Fm;* mm^3^); femur volume of the posterior leg *vs.* body surface ratio (*Fm:Bs;* mm^3^.mm^-2^); femur volume *vs.* tibia length of the posterior leg ratio (*Fm:Tb;* mm^3^.mm^-1^); membranous wing surface *vs.* body surface ratio (*Wg:Bs;* mm^2^.mm^-2^);
iii. *Resource acquisition-related traits:* mandible length *vs.* head surface ratio (*Md:Hd;* mm.mm^-2^); mandible length *vs.* labrum length ratio (*Md:Lb;* mm.mm^-1^).

All measurements were performed using a stereo-microscope (Leica Microsystems M50) equipped with an integrated high definition microscope camera (Leica IC80 HD).

### 2.4. Statistical analyses

#### Functional trait variations across carabid beetle species

We performed a principal component analysis (PCA) on the average traits of the 13 dominant species * 2 sexes to evaluate trait co-variations among species (Mouillot et al., 2013). We used a VARIMAX procedure to maximize correlations between PCA axes and traits. We then selected PCA axes with eigenvalue > 1 and recorded the PCA coordinates of each species. Then, for each species we calculated the mean values of each of the selected PCA axes that we used as species traits in the following analyses. This procedure has the advantage to select independent traits for analyses and help to define important leading dimensions of morphological variations between species (Maire et al., 2012).

An *a priori* hypothesis when using a mean trait value for each species is that intraspecific variability is sufficiently low so that the mean trait value of a species can be realistically used as a proxy for the species (Violle et al., 2012). To validate our approach, we thus estimated for each selected PCA axis the relative importance of intra and interspecific variability. In a linear model, we tested for the effect of species identity and sex on observed trait variability. Sex was nested within species. We then conducted a variance decomposition analysis based on the sum of square to estimate the importance of species and sex in explaining observed trait values. In a second analysis, we performed a linear discriminant analysis which finds the best linear combination of continuous explanatory variables (morphological traits of carabid beetles) separating different classes (here species) of a categorical variable. This analysis corresponds to another way to measure the importance of intraspecific trait variability (Albert et al., 2010).

#### Functional characterization of communities

We calculated the community-weighted mean (CWM) and variance (CWV) for each PCA axis separately:

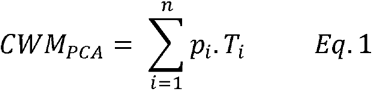

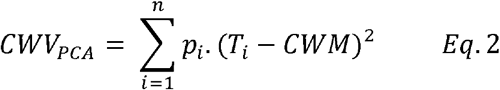

where *n* is the number of species in the community, *p_i_,* is the relative abundance of species *i* in a given community, *T_i_* is its PCA-based trait value. CWM_PCA_ reflects the mean PCA-based trait value of the community weighted by the abundance of each species (Violle et al., 2007). It reflects the functional identity of dominant species in a given community. CWV_PCA_ is a measurement of the functional diversity and quantifies the dispersion of PCA-based trait values within a given community (Le Bagousse□Pinguet et al., 2017). Calculated for each PCA-based trait separately, it is similar to commonly used distance-based indices of functional diversity such as functional dispersion or Rao index (Laliberté & Legendre, 2010).

#### Evaluation of the community assembly temporal dynamics

To investigate seasonal trends in community assembly and their variation between crops, linear mixed-effects models were run on CWM_PCA_s and CWV_PCA_s calculated on each selected PCA-based trait with the R package lme4 (Bates et al., 2015). CWM_PCA_ and CWV_PCA_ of each PCA-based trait were modelled separately as the response variable. To test whether carabid community assembly exhibited contrasted temporal trajectory in different crop types, we tested for an interaction between crop types (*Crop*) and time. Time was modelled as the scaled annual Julian date (*JD;* scaled with mean = 0). As carabid communities may be impacted by crop phenology or agricultural practices such as harvesting over the season (Marrec et al., 2015), we integrated in the model a polynomial order 3 for time to test for non-linear relationships. Sampling year (*Year*; n = 9) and preservative solution (*Solution;* n = 3) were included to account for sampling design. Field identity (*FieldID;* n = 625 levels) was included as a random intercept in all models, to account for within-year multi-sampling. Complete model formula was:

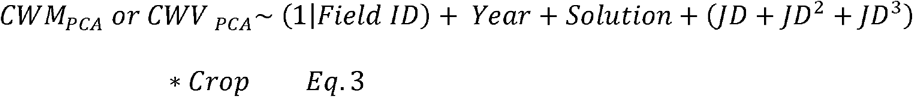

The best sub-model was selected by comparing AIC values (ΔAIC < 4) between all possible biologically relevant sub-models (*n* = 23) (Barton, 2013). Final model was estimated using restricted maximum likelihood (REML). To remove potential outliers, we excluded in prior analyses data points outside the upper quantile 99.9 % and then communities with less than three individuals (162 out of 1209 communities).

#### Evaluation of the impact of habitat filtering and niche differentiation

A null model approach was used to quantify the strength of PCA-based trait convergence and divergence in carabid beetle communities (CWV_PCA_) to isolate the impact of habitat filtering and niche differentiation (Götzenberger et al., 2016). The null hypothesis was that local communities should simply reflect a random distribution of individuals drawn from a regional species pool. As such, the regional species pool used to generate the null predictions must be carefully considered when inferring ecological processes from observed patterns (de Bello et al., 2012). As the regional species pool may vary over the season due to contrasted phenology between carabid beetles (Matalin, 2007) we constructed two alternative null models:

i. a global null model which considered the species pool observed throughout the growing season;
ii. a seasonal null model which took into account variations of carabid species pools over the growing season due to variation in phenology and agricultural practices between crop types.

A matrix describing the individual AA of each of the 13 species observed at the field scale was randomly shuffled 1,000 times across communities using the *permatful* function in the R package vegan (Oksanen et al., 2018). For the seasonal null model, the AA matrix was split according to three successive time periods (early spring: April 1 to May 15; late spring: May 15 to July 10; summer: July 10 to August 30; Fig. 1). Randomization was performed independently for each time period. Overall, the procedure kept species AA constant at the regional scale, but allowed species richness and AA to randomly vary between communities. Our individual-based randomization had the advantage to directly reflect our sampling design by taking into account the pattern of local AA of all sampled individuals at the community level. The size of the null envelope is only determined by species AA at the regional scale, consistently with our null hypothesis.

For each of the 1,000 randomizations and for the two null models, we used the matrix of trait values of each individual species to calculate the CWV_PCA_ at the community level. We then calculated the 95 % confidence interval to compare the observed CWV_PCA_ values to the predictions of the null model. If observed data felt outside of the null envelope, it indicated that deterministic processes led to less or more divergent community trait distribution than expected by chance. Specifically, observed CWV_PCA_ values below the null envelope indicated that traits within communities were forced to converge more than expected by chance, suggesting habitat filtering. In contrast, the impact of niche differentiation was detected when communities exhibited observed CWV_PCA_ values above the null envelope, i.e., coexisting species showed stronger functional differences than expected under the null hypothesis. As multiple assembly processes can simultaneously affect community structure and influence different traits independently (Gross et al., 2013; Spasojevic & Suding, 2012), we ran this analysis separately for each selected trait. For the seasonal null model, we also tested whether different crop types exhibited contrasted levels of trait dispersion (CWV_PCA_). To do so, we ran a linear mixed model such as described above for each period of time separately (early spring, late spring, and summer). The model had the following form:

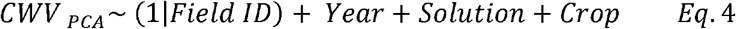

All statistical analyses were performed using the R environment (R. Core Team, 2018) and JMP11 (The SAS Institute, Cary, NC, USA).

## 3. RESULTS

### 3.1. Functional variations across carabid species

Body size, mobility, and resources acquisition traits defined three independent leading dimensions along which carabid species differentiated (total variance explained: 74 %; Fig. 2; see Table S3 in Appendix S2 for correlation tables). The first PCA axis (42 %) was associated to carabid body size and body shape (correlation with PCA axis for *Bs*: 0.93; *Bl:Bw:* 0.65), posterior leg size (*Lg:* 0.95; *Fm:Tb:* 0.51), femur size of the posterior leg (*Fm*: 0.95), and the relative proportion of their head surface and mandible length (*Md:Hd:* −0.87) (Fig. 2). The axis particularly opposed small species such as *Microlestes* spp. against large species such as *C. fuscipes* and *P. melanarius.*

**Fig. 2.**
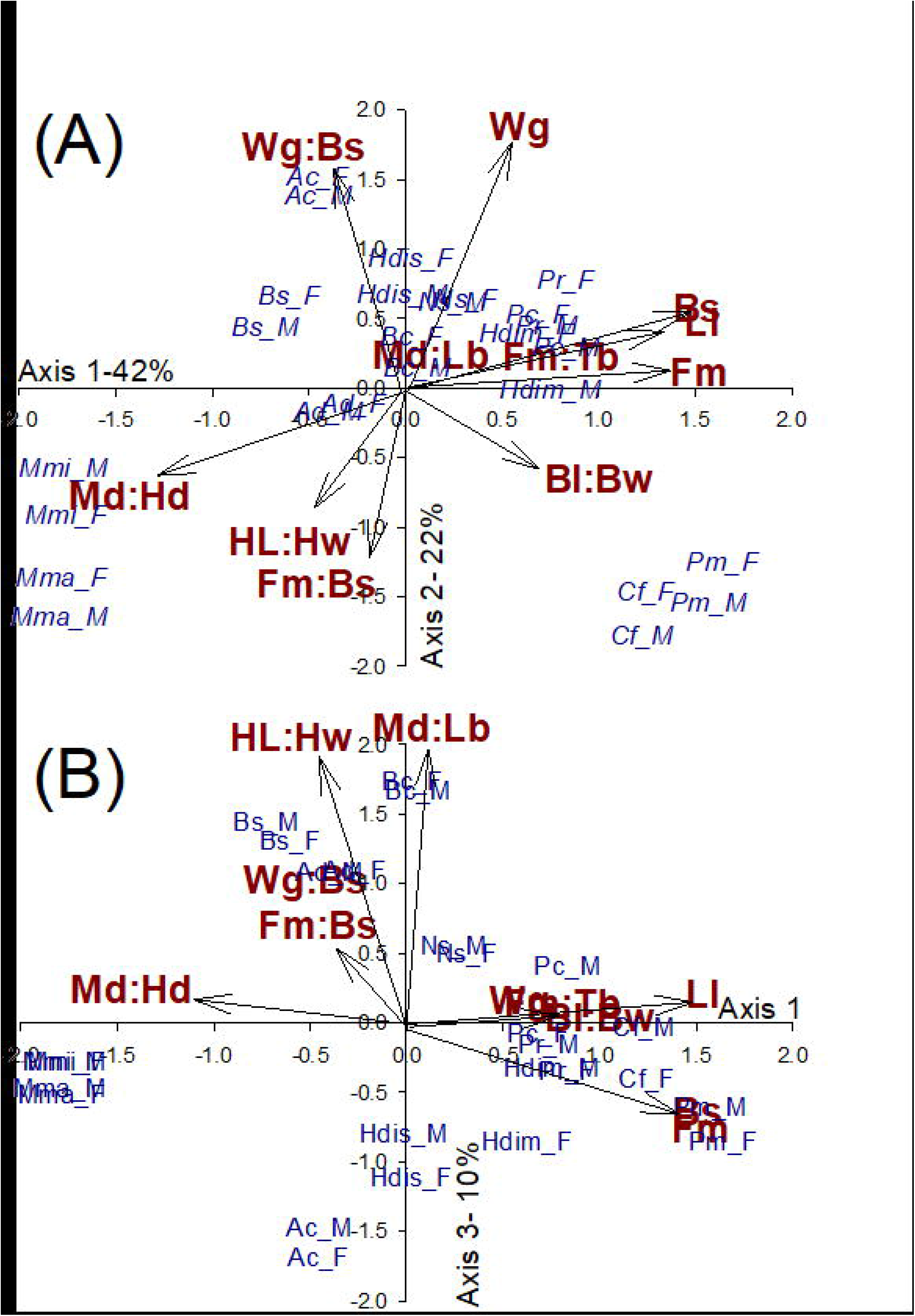
Co-variation of ground beetles’ morphological traits along the three first PCA axes (eigenvalue > 1). Morphological traits are in bold red, species are in blue. Trait abbreviations: *Bs,* body surface (mm^2^); *Lg,* leg length (mm); *Fm*, femur volume (mm^3^); *Wg,* membranous wing surface (mm^2^); *Bl:Bw,* body length *vs.* body width ratio (mm.mm^-1^); *Wg:Bs,* membranous wing surface *vs.* body surface ratio (mm^2^.mm^-2^); *Fm:Tb,* femur volume vs. tibia length ratio (mm^3^.mm^-1^); *Fm:Bs,* femur volume *vs.* body surface ratio (mm^3^.mm^-2^); *Md:Hd,* mandible length *vs.* head surface ratio (mm.mm^-2^); *Md:Lb,* mandible length *vs.* labrum length ratio (mm.mm^-1^); *Hl:Hw,* head length *vs.* head width (mm.mm^-1^). Species abbreviations are: Ac, *Amara consularis*; Ad, *Anchomenus dorsalis*; Bc, *Brachinus crepitans*; Bs, *Brachinus sclopeta*; Cf, *Calathus fuscipes;* Hdim, *Harpalus dimidiatus;* Hdis, *Harpalus distinguendus;* Mmi, *Microlestes minutulus;* Mma, *Microlestes maurus;* Ns, *Nebria salina;* Pc, *Poecilus cupreus;* Pr, *Pseudoophonus rufipes;* Pm, *Pterostichus melanarius.* M indicated male, F, female.

The second PCA axis (22 %) segregated species according to mobility traits and opposed species with large wings (*Wg:* 0.85; *Wg:Bs:* 0.74) to species with massive posterior legs (*Fm:Bs:* −0.67) and head larger than long (*Hl:Hw:* −0.56) (Fig. 2). The axis particularly opposed *A. consularis* and *H. distinguendus* against *M. maurus, C. fuscipes* and *P. melanarius*.

The third PCA axis (10 %) was mainly characterized by morphological traits linked to resource acquisition, and opposed species based on the relative length of their mandibles and labrum (*Md:Lb:* 0.59) and on the shape of their head (*Hl:Hw:* 0.56) (Fig. 2). The axis mainly opposed *Brachinus* spp. and *A. dorsalis* against *H. distinguendus* and *A. consularis.*

For the three PCA-based traits, intraspecific variability was very low compared to interspecific variability (%r^2^ = 0 to 3 %; Table 1). In addition, the linear discriminant analysis indicated only 11 % of misclassification due to intraspecific variability, confirming that interspecific variability was much stronger.

**Table 1.** Effect of interspecific differences and sexual dimorphism on trait variability. We indicated model parameter and proportion (%) of explained variance (%r^2^) by species and sex. We tested the effect of species and sex nested within species as explanatory variables and traits (PCA axes, see Fig. 2) as response variables.

| Traits | Adj. $r^2$ | Species | | | Sex | | |
| --- | --- | --- | --- | --- | --- | --- | --- |
| | | *Df* | *Fratio* | %$r^2$ | *Df* | *Fratio* | %$r^2$ |
| PCA axis 1 | 0.97 | 12 | 676.5 *** | 99.8 % | 13 | 1.4 | 0% |
| PCA axis 2 | 0.92 | 12 | 245.7 *** | 97.5 % | 13 | 6.3 *** | 3% |
| PCA axis 3 | 0.95 | 12 | 406.7 *** | 97.9 % | 13 | 8.3 *** | 2% |

### 3.2. Seasonal trends in the functional structure of carabid communities

For each response variable (hereafter named CWM_PCA1_, CWV_PCA1_, CWM_PCA2_, CWV_PCA2_, CWM_PCA3_, CWV_PCA3_ respectively for mean and dispersion of PCA axes 1, 2, and 3), 23 biologically relevant models were tested. The full model was the best or second best model for all response variables but CWM_PCA2_ (Appendix S3). CWM_PCA_ and CWV_PCA_ of all PCA-based traits varied significantly between crop types (except CWV_PCA3_, which was marginally significant) and community assembly significantly varied through the season, showing different trends among crops, for all PCA-based traits (Table 2).

**Table 2.** Effect of crop type and time on the functional structure of carabid communities. Values and significance of Type II Wald chi square tests realized on fixed effects selected in each of the ‘best’ (lower ΔAIC) final tested models after the selection procedure (see Appendix S3).

| Response variable | Fixed effects | Chisq | P (>Chisq) | Response variable | Fixed effects | Chisq | P (>Chisq) |
| --- | --- | --- | --- | --- | --- | --- | --- |
| **CWM1** | Year | 15.6 | < 0.001 | **CWV1** | Year | 30.8742 | < 0.001 |
|  | Solution | 18.6 | < 0.001 |  | Solution | 14.9308 | < 0.001 |
|  | JD | 2.89 | 0.089 |  | JD | 6.1195 | 0.0134 |
|  | JD<sup>2</sup> | 10.16 | 0.001 |  | JD<sup>2</sup> | 42.2013 | < 0.001 |
|  | JD<sup>3</sup> | 20.49 | < 0.001 |  | JD<sup>3</sup> | 8.3696 | 0.0038 |
|  | Crop | 84.57 | < 0.001 |  | Crop | 14.6505 | 0.0055 |
|  | JD:Crop | 43.23 | < 0.001 |  | JD:Crop | 15.2958 | 0.0041 |
|  | JD<sup>2</sup>:Crop | 31.15 | < 0.001 |  | JD<sup>2</sup>:Crop | - | - |
|  | JD<sup>3</sup>:Crop | - | - |  | JD<sup>3</sup>:Crop | 19.882 | < 0.001 |
| **CWM2** | Year | 2.11 | 0.146 | **CWV2** | Year | 13.652 | < 0.001 |
|  | Solution | 1.96 | 0.375 |  | Solution | 34.156 | < 0.001 |
|  | JD | 5.63 | 0.018 |  | JD | 50.93 | < 0.001 |
|  | JD<sup>2</sup> | 64.6 | < 0.001 |  | JD<sup>2</sup> | 29.081 | < 0.001 |
|  | JD<sup>3</sup> | 4.26 | 0.039 |  | JD<sup>3</sup> | 19.787 | < 0.001 |
|  | Crop | 29.27 | < 0.001 |  | Crop | 61.103 | < 0.001 |
|  | JD:Crop | - | - |  | JD:Crop | 20.486 | < 0.001 |
|  | JD<sup>2</sup>:Crop | 27.58 | < 0.001 |  | JD<sup>2</sup>:Crop | 18.572 | < 0.001 |
|  | JD<sup>3</sup>:Crop | - | - |  | JD<sup>3</sup>:Crop | 13.132 | 0.0107 |
| **CWM3** | Year | 0.05 | 0.826 | **CWV3** | Year | 0.86 | 0.355 |
|  | Solution | 3.84 | 0.146 |  | Solution | 1.86 | 0.395 |
|  | JD | 19.67 | < 0.001 |  | JD | 6.03 | 0.014 |
|  | JD<sup>2</sup> | 5.11 | 0.024 |  | JD<sup>2</sup> | 1.6 | 0.205 |
|  | JD<sup>3</sup> | 2.51 | 0.113 |  | JD<sup>3</sup> | 7.99 | 0.005 |
|  | Crop | 98.36 | < 0.001 |  | Crop | 9.14 | 0.058 |
|  | JD:Crop | 50.92 | < 0.001 |  | JD:Crop | 13.03 | 0.011 |
|  | JD<sup>2</sup>:Crop | 20.7 | < 0.001 |  | JD<sup>2</sup>:Crop | - | - |
|  | JD<sup>3</sup>:Crop | 21.26 | < 0.001 |  | JD<sup>3</sup>:Crop | 18.95 | < 0.001 |

For all PCA-based traits, CWM_PCA_s were not significantly different between crops at the very beginning and end of the growing season, but showed strong differences in their seasonal dynamic (Fig. 3). CWM_PCA1_ mainly linked to variation in body size, increased in sunflower to peak around June 9 and then decreased to the starting value (Fig. 3b). By contrast, CWM_PCA1_ decreased significantly in oilseed rape until ca. June 30 and then increased to the starting value (Fig. 3b). For the three other crops (alfalfa, grassland, and winter cereals), a gradual increase of CWM_PCA1_ was observed, to peak at the end of the season (Fig. 3a-b). For CWM_PCA2_, almost no temporal variation and differences between crops were observed, except for sunflower, in which it was lower than anywhere else in mid-spring, around ca. May 10-June 10, indicating communities mainly dominated by species with smaller wings and larger legs (Fig. 3d). For CWM_PCA3_, variations were mainly observed in annual crops, with higher values in oilseed rape than in other crops from ca. May 10 and which peaked around ca. June 30 and then decreased (Fig. 3f). This pattern tends to indicate than during this period, communities were dominated by species with relatively longer mandibles and heads. The exact opposite pattern was observed in sunflower during the same period (Fig. 3f). In winter cereals, CWM_PCA3_ was the highest around ca. April 30 and then decreased until the end of the season (Fig. 3f).

**Fig 3.**
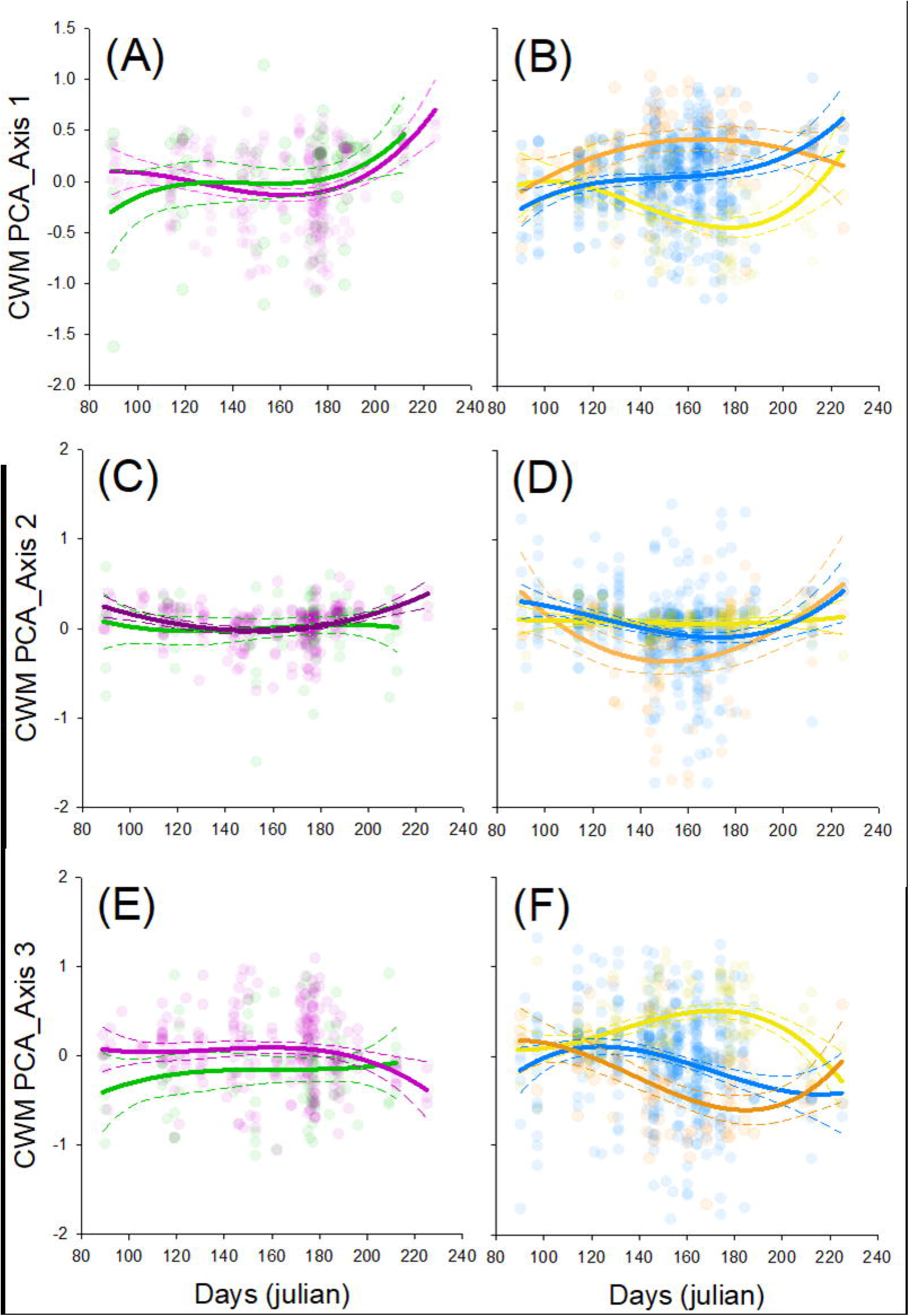
Change in community-weighted mean (CWM) through time (Julian days) for PCA axis 1, 2 and 3. We indicated model prediction for each crop (see Table 3 for model selection and parameters). In panels A, C and E we show model prediction for perennial crop, i.e. alfalfa (pink line) and grasslands (green line). In panels B, D and F we indicated model prediction for annual crops, i.e. wheat (blue line), oilseed rape (yellow line) and sunflower (orange line). Dots are raw data for each crop.

### 3.3. Evaluation of the impact of habitat filtering and niche differentiation

When considering the global null model, trait dispersion (CWV_PCA_) did not depart from the null envelop for any PCA-based trait and crop in the mid-season, except in sunflower for CWV_PCA2_ that was higher than expected by chance around ca. June 10-30 (Fig. 4d). CWV_PCA1_ and CWV_PCA2_ were lower than expected in alfalfa at the beginning of the season, and in winter cereals for CWV_PCA1_ at the end of the season (Fig. 4a-c). Functional diversity was not significantly higher in perennial crops than annual crops at any time, but tended to be higher in grassland at the very beginning of the season for CWV_PCA1_ (Fig. 4a).

**Fig 4.**
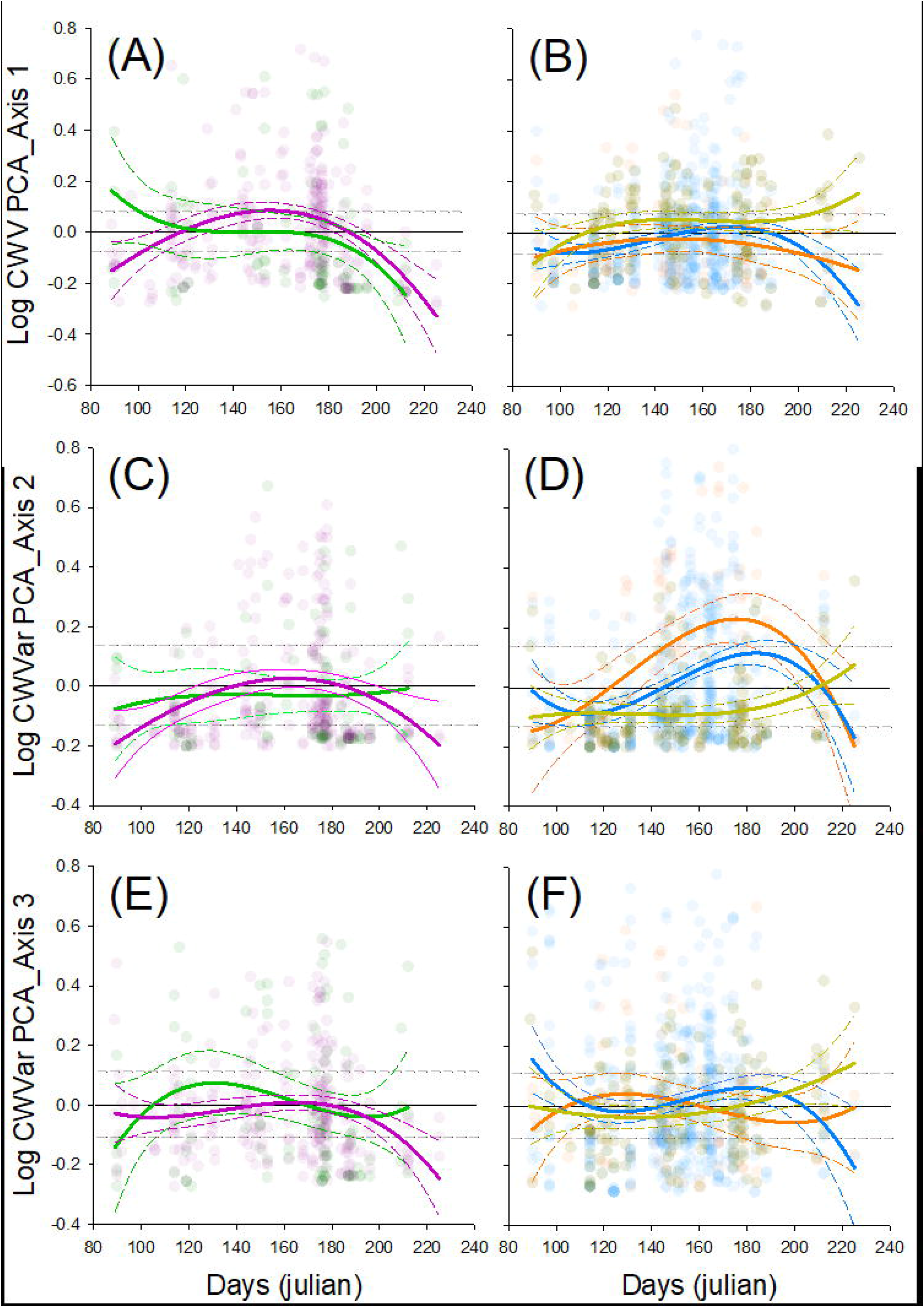
Change in community-weighted variance (log transformed, CWV) through time (Julian days) for PCA axis 1, 2 and 3. We indicated model prediction for each crop (see Table 3 for model selection and parameters). In panels A, C and E we show model prediction for perennial crop, i.e. alfalfa (pink line) and grasslands (green line). In panels B, D and F we indicated model prediction for annual crops, i.e. wheat (blue line), oilseed rape (yellow line) and sunflower (orange line). Dots are raw data. Predictions and dots were centered on the null model envelop (dash grey lines are the 95% confidence intervals).

When considering the seasonal null model, carabid community assembly highly changed through time for CWV_PCA1_ and CWV_PCA3_ while it was not the case for CWV_PCA2_ (Fig. 5). Community assembly did not significantly depart from the null envelop for any of the PCA-based traits in early spring, except for CWV_PCA1_ values which converged more than expected by chance in winter cereals. In late spring, CWV_PCA1_ and CWV_PCA3_ values were lower than expected by chance in all crops, suggesting trait convergence. Similar pattern was observed in summer, except in oilseed rape where values did not depart from the null expectation.

**Fig 5.**
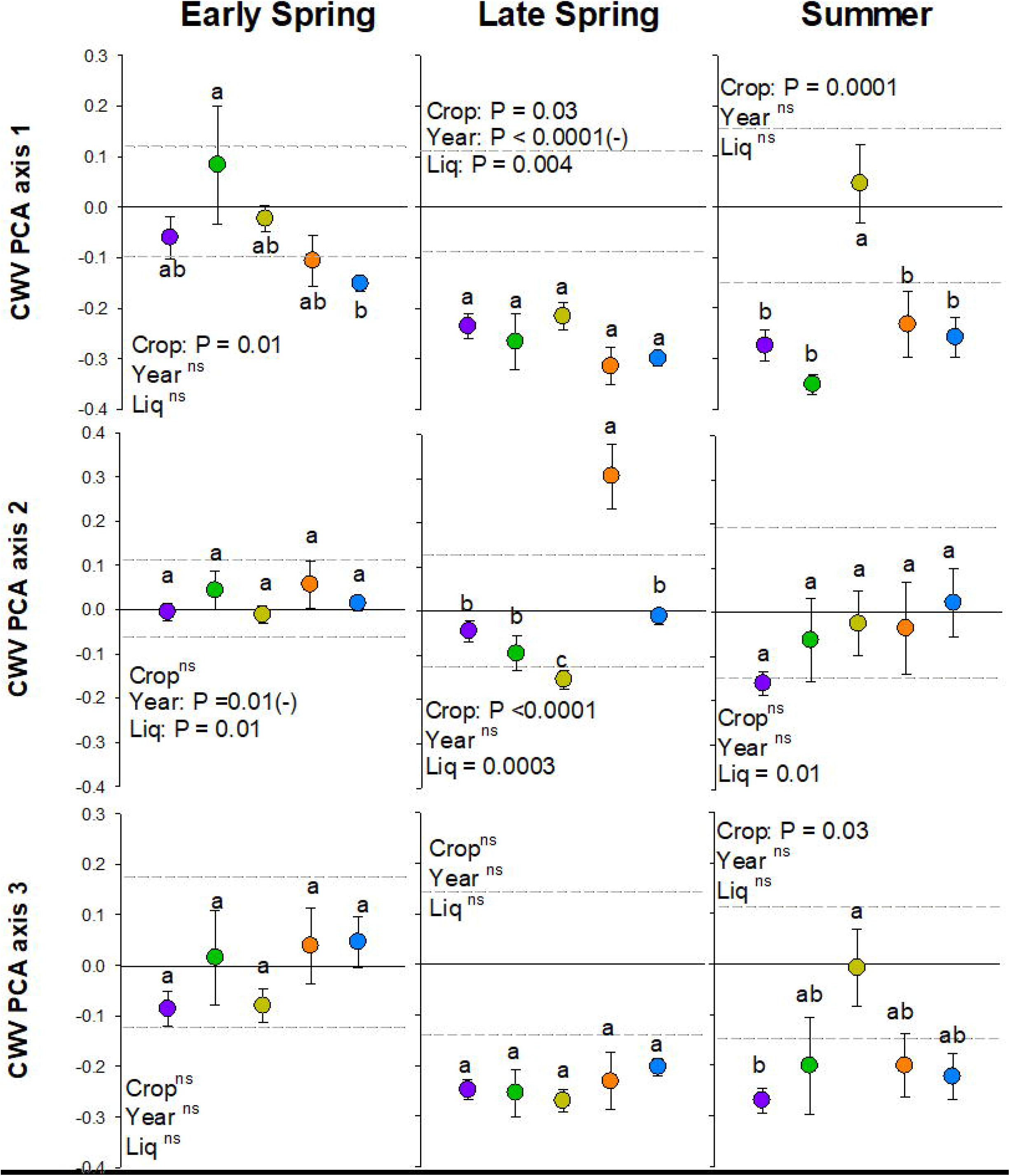
Community weighted variance (CWV) deviation from null prediction for traits 1, 2, and 3 in three successive time periods (early spring, April 1 to May 15; late spring, May15 to July 10); and summer July 10 to August 30). Each dot represents the observed CWV and associated standard error. Crops are: alfalfa (purple dots), grasslands (green dots), oilseed rape (yellow dots), sunflower (orange dots), and winter cereals (blue dots). Grey dashed lines are the 95% confidence interval of the null model envelop. We indicated for each CWV and periods model P values in each panel. Letters are *post hoc* Tukey HDS. For each panel, different letter indicated significant differences between crops.

## 4. DISCUSSION

In this multi-year study, we showed that the functional structure of carabid beetle communities varies importantly across the growing season in crops. This dynamic of community assembly for carabid beetles also strongly depends on the crop type. Although carabid beetles forms and functions widely vary across species, crops act as a habitat filter and strongly reduce the functional variation of co-occurring species within a field. However, we also showed that each crop type selects carabid species according to contrasting trait values and that dominant trait value could shift even within a single crop type over the season. Such high variability within and between crop types calls for the maintenance of diverse crop mosaics in agricultural landscapes (Sirami et al., 2019) to promote carabid species persistence, a key agent of biological pest control in agricultural landscapes.

### 4.1. Leading dimensions in functional traits across carabid species

Much functional ecology studies generally consider qualitative traits selected after a literature review and only partially available for all species. This generally impedes a precise characterization of all functional dimensions, because of unavailability of data for many species or trait variation across species distribution area. In our study, we measured continuous morphological traits on our sampled individuals. We found that carabid species, traits in arable field communities differentiate along three main dimensions of functional specialization, similar as previously shown for other taxa (Le Provost, Badenhausser, Le Bagousse-Pinguet, et al., 2020).

The first main dimension of interspecific differentiation was related to body size. Body size is associated with many life history traits and ecological characteristics that can explain its importance as a main driver of species assemblages. For instance, bigger carabid species have already been shown to be more prone to decline than smaller species when facing a loss of natural habitats, because of their lower reproductive rate and lower dispersal abilities (Kotze & O’Hara, 2003). Indeed, bigger species are expected to respond less rapidly to environmental changes than smaller species, which explains why communities found in farmlands are dominated by small and relatively unspecialized species (Aviron et al., 2005; Schweiger et al., 2005).

The second leading dimension was based on mobility traits. Species appear to importantly oppose according to whether they have larger, well developed wings, or larger and stronger legs. Carabid beetles exhibit a variety of wing attributes, including wing dimorphism, which can have implications for their dispersal abilities (Kotze et al., 2011). However, the shape of posterior legs is also correlated, in carabid beetles and other Coleoptera, to movement ability and ecological behaviour, especially speed attained and pushing force (Evans & Forsythe, 1984). Relatively short legs or/and short and slender femora are expected in horizontal pusher species, with reduced movement abilities, in opposition with species with relatively long legs and large femora which are faster runners but weak pushers (Evans & Forsythe, 1984; Forsythe, 1983).

The last dimension of interspecific differentiation was based on resource-acquisition traits. An opposition appears between species with relatively longer head or/and mandibles and species with relatively broader head or/and shorter mandibles. Previous studies (e.g., Acorn & Ball, 1991; Deroulers & Bretagnolle, 2019; Kulkarni et al., 2015) correlated a phytophagous diet to more robust, broader mandibles in carabids, which is in accordance with feeding niche information obtained from the literature for our species (Appendix S4). However, information about carabid diet is relatively unknown for most species, and current knowledge is often based on individual observations or lab experiments (Deroulers & Bretagnolle, 2019). Better evaluation of carabid diet is required, using alternative approaches, such as gut content analysis (Kamenova et al., 2018) or isotopic and fatty acid composition analysis (González Macé et al., 2019).

### 4.2. Habitat filtering shapes carabid communities in crops

We find support for one of our research hypotheses: each crop type acts as a habitat filter on carabid beetles, filtering out species according to their functional trait values, when taking the seasonal variation of the species pool into account (hypothesis 2). Functional diversity was on average lower than expected by chance under a random assembly of local communities. Nonetheless, the strength of this pattern varied depending on the null model considered. When considering a global null model based on the entire species pool observed throughout the growing season, communities seem randomly assembled or even subject to niche differentiation processes in late spring, which could have led to an erroneous validation of hypothesis 1. In fact, this period corresponds in temperate regions to a transition in carabid community composition. In early spring, communities are composed of “spring breeders” (Thiele, 1977). From mid-spring, there is an increase of the regional species pool due to the emergence of “summer-autumn breeders” (Matalin, 2008). As a consequence, there is a sudden increase of the functional diversity in local assemblages which can falsely be interpreted as niche differentiation processes operating at the field scale. On the contrary, when considering a seasonal null model, which takes into account variations of carabid species pools over the growing season, a clear habitat filtering pattern was revealed. To sum up, we showed that whether or not integrating temporal change in species pool when investigating functional assembly dynamics can lead to very different interpretations and conclusions. Although carabid communities may show strong patterns of temporal niche differentiation (e.g., in forest ecosystems, Loreau, 1989), especially through competition processes (Kamenova et al., 2015), they are generally strongly filtered within fields in response to crop type and crop phenology.

Habitat filtering was observed in all crop types, indicating high specialization of carabid communities at the crop level, as previously suggested (Marrec et al., 2015; Weibull & Östman, 2003). Crop habitat filtering was the highest in late spring, while it was almost inexistent in early spring. This seemingly random assembly of species in early spring can be explained by the fact that abundance-activity of carabid species in fields does not entirely depend on the present crop type, but mainly on the previous crop type(s) in the succession, and on the landscape context as carabid may colonize crops from nearby habitats at the onset of the growing season (e.g., Marrec et al., 2015, 2017). Because of crop rotation, farmlands are highly dynamic landscapes, both in space and time. To face induced brutal changeovers, carabid individuals of most species might have to redistribute between fields of different crop types, and between crops and non-crop habitats before winter to shelter for overwintering, and in early spring to find a new suitable habitat patch (Geiger et al., 2009; Holland et al., 2005; Marrec et al., 2015; Thomas et al., 2001). Similar distribution shifts of individuals among crops or/and non-crop habitats have also been reported in summer, when spring-summer crops become more attractive as they grow, while winter crops are ripening, drying, and then harvested (O’Rourke et al., 2014; Schneider et al., 2016). Similarly as in early spring, these summer distribution shifts may explain the lower habitat filtering we found in summer.

Crop types did not select the same trait values, and we observed high functional specializations. Carabid species distribution depends mainly on microclimatic conditions and availability of resources (Lövei & Sunderland, 1996), which differ importantly between crops, due to differences in crop practices, crop phenology, vegetation structure, etc. Body size and resource-acquisition traits were the most affected by crop type in all crops. In late spring, oilseed rape species assemblages were characterized by small species with relatively long mandibles, traits associated to small predators (such as *M. <u>maurus</u>, M. minutulus,* and *B. sclopeta).* Oilseed rape fields are generally highly affected by many pest species, and their understory moisture conditions shelter many arthropod species, which can be as many potential preys for predators (e.g., Zaller et al., 2009). The reverse pattern was observed in sunflower, with larger carabid species with shorter mandibles, more characteristic of phytophagous diets (such as *H. dimidiatus* and *H. distinguendus).* Sunflower fields are sown in April in our study area (Fig. 1), which means soils are disturbed in early spring, and vegetative ground cover and pest species abundance are still low in late spring. As a consequence, phytophagous species generally dominate carabid assemblages in more disturbed habitats (Ribera et al., 2001). Consistently, soil ploughing allows buried seeds to resurface, and then provide food resources for granivorous species. In a recent study (Labruyère et al., 2016), AA of generalist phytophagous and polyphagous carabid species was congruently shown to be higher in spring crops (sugar beet, maize, and spring oilseed rape) than in winter oilseed rape. Finally, intermediate morphologies are found in all other crops (winter cereals, alfalfa, and grassland). Higher medium-sized beetle activity has already been shown in grassy habitats compared to annual crops (Eyre et al., 2009), with body size decreasing in more intensively managed habitats (Blake et al., 1994).

On the other hand, our results did not show strong selection for species in considered crop types depending on mobility attributes, at any time. The first reason would be that the ability to disperse is likely to be selected at a scale much larger than the field: the landscape scale. Dispersal-related traits might be filtered by landscape spatiotemporal structure. It has been previously shown that long-term land-use change to more intensive agricultural landscapes has impoverished the functional diversity of mobility traits in carabid assemblages (Le Provost, Badenhausser, Le Bagousse-Pinguet, et al., 2020), and selected species with higher dispersal abilities and tolerance against agricultural disturbances (Turin & Den Boer, 1988). Secondly, in some species, macropterous individuals do not necessarily possess functioning flight muscles and are therefore incapable of flight (Desender & Turin, 1989; Nelemans, 1987), at least at certain periods of their life cycle (Van Huizen, 1977). Such an evaluation is arduous, especially in smaller species, but would allow to better understanding intra and interspecific variations between flying and walking strategies to reach new habitat patches.

Recent studies have highlighted the importance of crop diversity in the landscape to maintain diverse arthropod communities in farmlands (e.g., Fahrig et al., 2011; Sirami et al., 2019). Two distinct hypotheses have been proposed to explain the effect of crop diversity: crop diversity should benefit biodiversity if many species are either specialist of distinct crop types (i.e., habitat specialization; Weibull et al., 2003) or require multiple resources provided by different crop types (i.e., landscape complementation; Dunning et al., 1992). In our study, we showed that different crops host contrasted carabid communities as crop type filters specific trait values. Such strong functional specialization of carabid communities may explain the effect of crop diversity that has been observed across many agricultural regions (including ours, see Sirami et al., 2019) and supports the habitat specialization hypothesis. However, by focusing our study on seasonal changes observed over the spring-summer period, we also show that contrasted carabid communities can inhabit a same crop type but at different periods of the year. Therefore, our results also support the landscape complementation hypothesis operating over time on carabid communities.

## 5. CONCLUSIONS

Most studies on carabid beetles functional traits have focused on size-related traits or literature-based information. Recent studies have proposed that morphological diversification of carabid beetles strongly respond to environmental heterogeneity and help to identify contrasted functional specialization (other than size) axes by determining their response to increasing environmental pressure (Le Provost, Badenhausser, Le Bagousse-Pinguet, et al., 2020). Combining various morphological traits related to different ecological functions, we found support to our second hypothesis: each crop type acts as a filter on carabid communities for body size and resource-acquisition traits, and functional differentiation between crops increases with time during crop growing season. Such variations seem to be mainly related to crop type and phenology. However, other drivers such as disturbances involved by agricultural practices associated with crop types may also explain the observed patterns. As highly mobile organisms, carabid beetles are able to follow high temporal fluctuations of crop distribution and resources. As such, mobility traits were not selected by crop types, but more likely at the landscape scale. Indeed, intensive agricultural landscapes are dominated by crops with short rotation times even in the case of grasslands which are mainly temporary grasslands included in crop rotations. Consequently, because of the strong functional specialisation to habitat type, maintaining high diversity of crops and non-crop habitats seems crucial to sustain carabid communities in agroecosystems.

## ACKNOWLEDGEMENTS

We are especially grateful to the numerous field workers who helped collect the data. We also thank all involved farmers for letting us work on their land. R.M. was funded by a Région Poitou-Charentes and Département des Deux-Sèvres Ph.D. grant. This work is part of the Landscaphid project and was supported by the ANR Systerra program (French National Research agency, ANR-09-STRA-05) and by the French national DIVA2 program. Authors declare to not having any conflict of interest.

## AUTHORS, CONTRIBUTIONS

IB, VB, GC, BG, NG, and RM designed the study.

BG, NG, and RM wrote the manuscript.

BG, AD, RM, and MR collected the data.

MR sorted arthropods from the traps and identified the species.

AD measured the morphological traits.

AD and NG ran the statistical analyses.

All authors contributed critically to the drafts and gave final approval for publication.

## REFERENCES

Acorn, J. H., & Ball, G. E. (1991). The mandibles of some adult ground beetles: Structure, function, and the evolution of herbivory (Coleoptera: Carabidae). Canadian Journal of Zoology, 69(3), 638–650.

Albert, C. H., Thuiller, W., Yoccoz, N. G., Douzet, R., Aubert, S., & Lavorel, S. (2010). A multi-trait approach reveals the structure and the relative importance of intra-vs. Interspecific variability in plant traits. Functional Ecology, 24(6), 1192–1201.

Aviron, S., Burel, F., Baudry, J., & Schermann, N. (2005). Carabid assemblages in agricultural landscapes: Impacts of habitat features, landscape context at different spatial scales and farming intensity. Agriculture, Ecosystems & Environment, 108(3), 205–217.

Barton, K. (2013). MuMIn: Multi-model inference, R package version 1.9.0. http://R-Forge.r-Project.Org/Projects/Mumin/.

Bates, D., Maechler, M., Bolker, B., & Walker, S. (2015). Fitting Linear Mixed-Effects Models Using Ime4. Journal of Statistical Software, 67(1), 1–48. https://doi.org/doi:10.18637/jss.v067.i01

Blake, S., Foster, G. N., Eyre, M. D., & Luff, M. L. (1994). Effects of habitat type and grassland management practices on the body size distribution of carabid beetles. Pedobiologia, 38(6), 502–512.

Bretagnolle, V., Berthet, E., Gross, N., Gauffre, B., Plumejeaud, C., Houte, S., Badenhausser, I., Monceau, K., Fabrice, A., & Monestiez, P. (2018). Biodiversity, ecosystem services and citizen science: The value of long term monitoring in farmland landscapes for sustainable agriculture. Sci. Total Environ, 627, 822–834.

Bretagnolle, Vincent, Berthet, E., Gross, N., Gauffre, B., Plumejeaud, C., Houte, S., Badenhausser, I., Monceau, K., Allier, F., Monestiez, P., & Gaba, S. (2018). Description of long-term monitoring of farmland biodiversity in a LTSER. Data in Brief, 19, 1310–1313. https://doi.org/10.1016/j.dib.2018.05.028

Brown, J. H., Gillooly, J. F., Allen, A. P., Savage, V. M., & West, G. B. (2004). Toward a metabolic theory of ecology. Ecology, 85(7), 1771–1789.

Chesson, P. (2000). Mechanisms of maintenance of species diversity. Annual Review of Ecology and Systematics, 31(1), 343–366.

de Bello, F., Price, J. N., Münkemüller, T., Liira, J., Zobel, M., Thuiller, W., Gerhold, P., Götzenberger, L., Lavergne, S., & Lepš, J. (2012). Functional species pool framework to test for biotic effects on community assembly. Ecology, 93(10), 2263–2273.

de Jong, Y., Verbeek, M., Michelsen, V., de Place Bjørn, P., Los, W., Steeman, F., Bailly, N., Basire, C., Chylarecki, P., & Stloukal, E. (2014). Fauna Europaea–all European animal species on the web. Biodiversity Data Journal, 2, e4034. https://doi.org/10.3897/BDJ.2.e4034

Deroulers, P., & Bretagnolle, V. (2019). The consumption pattern of 28 species of carabid beetles (Carabidae) to a weed seed, Viola arvensis. Bulletin of Entomological Research, 109(2), 229–235.

Desender, K., & Turin, H. (1989). Loss of habitats and changes in the composition of the ground and tiger beetle fauna in four West European countries since 1950 (Coleoptera: Carabidae, Cicindelidae). Biological Conservation, 48(4), 277–294. https://doi.org/10.1016/0006-3207(89)90103-1

Dunning, J. B., Danielson, B. J., & Pulliam, H. R. (1992). Ecological processes that affect populations in complex landscapes. Oikos, 169–175.

Evans, M. E. G., & Forsythe, T. G. (1984). A comparison of adaptations to running, pushing and burrowing in some adult Coleoptera: Especially Carabidae. Journal of Zoology, 202(4), 513–534.

Eyre, M. D., Labanowska-Bury, D., Avayanos, J. G., White, R., & Leifert, C. (2009). Ground beetles (Coleoptera, Carabidae) in an intensively managed vegetable crop landscape in eastern England. Agriculture, Ecosystems & Environment, 131(3–4), 340–346.

Eyre, M. D., Luff, M. L., & Leifert, C. (2013). Crop, field boundary, productivity and disturbance influences on ground beetles (Coleoptera, Carabidae) in the agroecosystem. Agriculture, Ecosystems & Environment, 165, 60–67.

Fahrig, L., Baudry, J., Brotons, L., Burel, F. G., Crist, T. O., Fuller, R. J., Sirami, C., Siriwardena, G. M., & Martin, J.-L. (2011). Functional landscape heterogeneity and animal biodiversity in agricultural landscapes. Ecology Letters, 14(2), 101–112.

Fitzgerald, D. B., Winemiller, K. O., Sabaj Pérez, M. H., & Sousa, L. M. (2017). Seasonal changes in the assembly mechanisms structuring tropical fish communities. Ecology, 98(1), 21–31.

Forsythe, T. G. (1983). Locomotion in ground beetles (Coleoptera Carabidae): An interpretation of leg structure in functional terms. Journal of Zoology, 200(4), 493–507.

Fukami, T., Bezemer, T. M., Mortimer, S. R., & van der Putten, W. H. (2005). Species divergence and trait convergence in experimental plant community assembly. Ecology Letters, 8(12), 1283–1290.

Gaüzère, P., Jiguet, F., & Devictor, V. (2015). Rapid adjustment of bird community compositions to local climatic variations and its functional consequences. Global Change Biology, 21(9), 3367–3378.

Geiger, F., Wäckers, F. L., & Bianchi, F. J. (2009). Hibernation of predatory arthropods in semi-natural habitats. BioControl, 54(4), 529–535.

González Macé, O., Ebeling, A., Eisenhauer, N., Cesarz, S., & Scheu, S. (2019). Variations in trophic niches of generalist predators with plant community composition as indicated by stable isotopes and fatty acids. SOIL ORGANISMS, 91(2), 45–59–45–59.

Götzenberger, L., Botta-Dukát, Z., Lepš, J., Pärtel, M., Zobel, M., & de Bello, F. (2016). Which randomizations detect convergence and divergence in trait-based community assembly? A test of commonly used null models. Journal of Vegetation Science, 27(6), 1275–1287.

Grime, J. P. (2006). Trait convergence and trait divergence in herbaceous plant communities: Mechanisms and consequences. Journal of Vegetation Science, 17(2), 255–260.

Gross, N., Börger, L., Soriano-Morales, S. I., Le Bagousse-Pinguet, Y., Quero, J. L., García-Gómez, M., Valencia-Gómez, E., & Maestre, F. T. (2013). Uncovering multiscale effects of aridity and biotic interactions on the functional structure of Mediterranean shrublands. Journal of Ecology, 101(3), 637–649.

Gross, N., Liancourt, P., Butters, R., Duncan, R. P., & Hulme, P. E. (2015). Functional equivalence, competitive hierarchy and facilitation determine species coexistence in highly invaded grasslands. New Phytologist, 206(1), 175–186.

Habel, J. C., Seibold, S., Ulrich, W., & Schmitt, T. (2018). Seasonality overrides differences in butterfly species composition between natural and anthropogenic forest habitats. Animal Conservation.

HilleRisLambers, J., Adler, P. B., Harpole, W. S., Levine, J. M., & Mayfield, M. M. (2012). Rethinking community assembly through the lens of coexistence theory. Annual Review of Ecology, Evolution, and Systematics, 43, 227–248.

Holland, J. M., Birkett, T., & Southway, S. (2009). Contrasting the farm-scale spatio-temporal dynamics of boundary and field overwintering predatory beetles in arable crops. Biocontrol, 54(1), 19–33.

Holland, J. M., Thomas, C. F. G., Birkett, T., Southway, S., & Oaten, H. (2005). Farm-scale spatiotemporal dynamics of predatory beetles in arable crops. Journal of Applied Ecology, 42(6), 1140–1152.

Hubbell, S. P. (2005). Neutral theory in community ecology and the hypothesis of functional equivalence. Functional Ecology, 19(1), 166–172.

Jeannel, R. (1941). Coléoptères Carabiques, Faune de France. Lechevalier.

Jeannel, R. (1942). Coléoptères Carabiques II, Faune de France. Lechevalier.

Kamenova, S., Mayer, R., Rubbmark, O. R., Coissac, E., Plantegenest, M., & Traugott, M. (2018). Comparing three types of dietary samples for prey DNA decay in an insect generalist predator. Molecular Ecology Resources, 18(5), 966–973.

Kamenova, S., Tougeron, K., Cateine, M., Marie, A., & Plantegenest, M. (2015). Behaviour-driven micro-scale niche differentiation in carabid beetles. Entomologia Experimentalis et Applicata, 155(1), 39–46.

Keddy, P. A. (1992). Assembly and response rules: Two goals for predictive community ecology. Journal of Vegetation Science, 3(2), 157–164.

Kotze, D. J., Brandmayr, P., Casale, A., Dauffy-Richard, E., Dekoninck, W., Koivula, M. J., Lövei, G. L., Mossakowski, D., Noordijk, J., & Paarmann, W. (2011). Forty years of carabid beetle research in Europe–from taxonomy, biology, ecology and population studies to bioindication, habitat assessment and conservation. ZooKeys, 100, 55.

Kotze, D. J., & O’Hara, R. B. (2003). Species decline—But why? Explanations of carabid beetle (Coleoptera, Carabidae) declines in Europe. Oecologia, 135(1), 138–148.

Kraft, N. J., Valencia, R., & Ackerly, D. D. (2008). Functional traits and niche-based tree community assembly in an Amazonian forest. Science, 322(5901), 580–582.

Kromp, B. (1999). Carabid beetles in sustainable agriculture: A review on pest control efficacy, cultivation impacts and enhancement. In Invertebrate biodiversity as bioindicators of sustainable landscapes (pp. 187–228). Elsevier.

Kulkarni, S. S., Dosdall, L. M., & Willenborg, C. J. (2015). The role of ground beetles (Coleoptera: Carabidae) in weed seed consumption: A review. Weed Science, 63(2), 355–376.

Labruyère, S., Bohan, D. A., Biju-Duval, L., Ricci, B., & Petit, S. (2016). Local, neighbor and landscape effects on the abundance of weed seed-eating carabids in arable fields: A nationwide analysis. Basic and Applied Ecology, 17(3), 230–239.

Laliberté, E., & Legendre, P. (2010). A distance-based framework for measuring functional diversity from multiple traits. Ecology, 91(1), 299–305.

Le Bagousse-Pinguet, Y., Gross, N., Maestre, F. T., Maire, V., De Bello, F., Fonseca, C. R., Kattge, J., Valencia, E., Leps, J., & Liancourt, P. (2017). Testing the environmental filtering concept in global drylands. Journal of Ecology, 105(4), 1058–1069.

Le Provost, G., Badenhausser, I., Le Bagousse-Pinguet, Y., Clough, Y., Henckel, L., Violle, C., Bretagnolle, V., Roncoroni, M., Manning, P., & Gross, N. (2020). Land-use history impacts functional diversity across multiple trophic groups. Proceedings of the National Academy of Sciences.

Le Provost, G., Badenhausser, I., Violle, C., Requier, F., D’Ottavio, M., Roncoroni, M., Gross, L., & Gross, N. (2020). Grassland-to-crop conversion in agricultural landscapes has lasting impact on the trait diversity of bees. Landscape Ecology, 1–15.

Le Provost, G., Gross, N., Börger, L., Deraison, H., Roncoroni, M., & Badenhausser, I. (2017). Trait-matching and mass effect determine the functional response of herbivore communities to land-use intensification. Functional Ecology, 31(8), 1600–1611.

Loreau, M. (1989). On testing temporal niche differentiation in carabid beetles. Oecologia, 81(1), 89–96.

Lövei, G. L., & Sunderland, K. D. (1996). Ecology and behavior of ground beetles (Coleoptera: Carabidae). Annual Review of Entomology, 41(1), 231–256.

MacArthur, R., & Levins, R. (1967). The limiting similarity, convergence, and divergence of coexisting species. The American Naturalist, 101(921), 377–385.

Maire, V., Gross, N., Börger, L., Proulx, R., Wirth, C., Pontes, L. da S., Soussana, J.-F., & Louault, F. (2012). Habitat filtering and niche differentiation jointly explain species relative abundance within grassland communities along fertility and disturbance gradients. New Phytologist, 196(2), 497–509.

Marrec, R., Badenhausser, I., Bretagnolle, V., Börger, L., Roncoroni, M., Guillon, N., & Gauffre, B. (2015). Crop succession and habitat preferences drive the distribution and abundance of carabid beetles in an agricultural landscape. Agriculture, Ecosystems & Environment, 199, 282–289.

Marrec, R., Caro, G., Miguet, P., Badenhausser, I., Plantegenest, M., Vialatte, A., Bretagnolle, V., & Gauffre, B. (2017). Spatiotemporal dynamics of the agricultural landscape mosaic drives distribution and abundance of dominant carabid beetles. Landscape Ecology, 32(12), 2383–2398.

Matalin, A. V. (2007). Typology of life cycles of ground beetles (Coleoptera, Carabidae) in Western Palaearctic. Entomological Review, 87(8), 947–972.

Matalin, Andrey V. (2008). Evolution of biennial life cycles in ground beetles (Coleoptera, Carabidae) of the Western Palaearctic. Back to the Roots and Back to the Future. Proceedings of XIII European Carabidologist Meeting, Blagoevgrad, 259–284.

Moretti, M., Dias, A. T., De Bello, F., Altermatt, F., Chown, S. L., Azcárate, F. M., Bell, J. R., Fournier, B., Hedde, M., & Hortal, J. (2017). Handbook of protocols for standardized measurement of terrestrial invertebrate functional traits. Functional Ecology, 31(3), 558–567.

Mouillot, D., Graham, N. A., Villéger, S., Mason, N. W., & Bellwood, D. R. (2013). A functional approach reveals community responses to disturbances. Trends in Ecology & Evolution, 28(3), 167–177.

Mouquet, N., Munguia, P., Kneitel, J. M., & Miller, T. E. (2003). Community assembly time and the relationship between local and regional species richness. Oikos, 103(3), 618–626.

Nelemans, M. N. E. (1987). Possibilities for flight in the carabid beetle Nebria brevicollis (F.). Oecologia, 72(4), 502–509. https://doi.org/10.1007/BF00378974

Newbold, T., Hudson, L. N., Hill, S. L., Contu, S., Lysenko, I., Senior, R. A., Börger, L., Bennett, D. J., Choimes, A., & Collen, B. (2015). Global effects of land use on local terrestrial biodiversity. Nature, 520(7545), 45.

Oksanen, J., Blanchet, F. G., Friendly, M., Kindt, R., Legendre, P., McGlinn, D., Minchin, P. R., O’Hara, R. B., Simpson, G. L., Solymos, P., Stevens, H. H., Szoecs, E., & Wagner, H. (2018). Vegan: Community Ecology Package. R package version 2.5-2. https://CRAN.R-project.org/package=vegan.

O’Rourke, M. E., Liebman, M., & Rice, M. E. (2014). Ground beetle (Coleoptera: Carabidae) assemblages in conventional and diversified crop rotation systems. Environmental Entomology, 37(1), 121–130.

Pakeman, R. J., & Stockan, J. A. (2014). Drivers of carabid functional diversity: Abiotic environment, plant functional traits, or plant functional diversity? Ecology, 95(5), 1213–1224.

R. Core Team. (2018). R: A language and environment for statistical computing. R Foundation for Statistical Computing, Vienna, Austria. URL https://www.R-project.org/.

Ribera, L., Dolédec, S., Downie, I. S., & Foster, G. N. (2001). Effect of land disturbance and stress on species traits of ground beetle assemblages. Ecology, 82(4), 1112–1129.

Rusch, A., Birkhofer, K., Bommarco, R., Smith, H. G., & Ekbom, B. (2015). Predator body sizes and habitat preferences predict predation rates in an agroecosystem. Basic and Applied Ecology, 16(3), 250–259.

Scheffer, M., Carpenter, S., Foley, J. A., Folke, C., & Walker, B. (2001). Catastrophic shifts in ecosystems. Nature, 413(6856), 591.

Schneider, G., Krauss, J., Boetzl, F. A., Fritze, M.-A., & Steffan-Dewenter, I. (2016). Spillover from adjacent crop and forest habitats shapes carabid beetle assemblages in fragmented semi-natural grasslands. Oecologia, 182(4), 1141–1150.

Schweiger, O., Maelfait, J.-P., Van Wingerden, W., Hendrickx, F., Billeter, R., Speelmans, M., Augenstein, I., Aukema, B., Aviron, S., & Bailey, D. (2005). Quantifying the impact of environmental factors on arthropod communities in agricultural landscapes across organizational levels and spatial scales. Journal of Applied Ecology, 42(6), 1129–1139.

Sirami, C., Gross, N., Baillod, A. B., Bertrand, C., Carrié, R., Hass, A., Henckel, L., Miguet, P., Vuillot, C., & Alignier, A. (2019). Increasing crop heterogeneity enhances multitrophic diversity across agricultural regions. Proceedings of the National Academy of Sciences, 116(33), 16442–16447.

Skvarla, M. J., Larson, J. L., & Dowling, A. P. G. (2014). Pitfalls and preservatives: A review. The Journal of the Entomological Society of Ontario, 145, 15–43.

Spasojevic, M. J., & Suding, K. N. (2012). Inferring community assembly mechanisms from functional diversity patterns: The importance of multiple assembly processes. Journal of Ecology, 100(3), 652–661.

Thiele, H.-U. (1977). Carabid Beetles in Their Environments. A Study on Habitat Selection by Adaptations in Physiology and Behaviour. Springer-Verlag.

Thomas, C. F. G., Parkinson, L., Griffiths, G. J. K., Garcia, A. F., & Marshall, E. J. P. (2001). Aggregation and temporal stability of carabid beetle distributions in field and hedgerow habitats. Journal of Applied Ecology, 38(1), 100–116.

Turin, H., & Den Boer, P. J. (1988). Changes in the distribution of carabid beetles in The Netherlands since 1880. II. Isolation of habitats and long-term time trends in the occurence of carabid species with different powers of dispersal (Coleoptera, Carabidae). Biological Conservation, 44(3), 179–200.

Uvarov, B. (1977). Grasshoppers and Locusts. A Handbook of General Acridology Vol. 2. Behaviour, Ecology, Biogeography, Population Dynamics. Centre for Overseas Pest Research.

Van Huizen, T. H. P. (1977). The significance of flight activity in the life cycle of Amara plebeja Gyll.(Coleoptera, Carabidae). Oecologia, 29(1), 27–41.

Vanneste, T., Valdés, A., Verheyen, K., Perring, M. P., Bernhardt-Römermann, M., Andrieu, E., Brunet, J., Cousins, S. A., Deconchat, M., & De Smedt, P. (2019). Functional trait variation of forest understorey plant communities across Europe. Basic and Applied Ecology, 34, 1–14.

Violle, C., Enquist, B. J., McGill, B. J., Jiang, L. I. N., Albert, C. H., Hulshof, C., Jung, V., & Messier, J. (2012). The return of the variance: Intraspecific variability in community ecology. Trends in Ecology & Evolution, 27(4), 244–252.

Violle, C., Navas, M.-L., Vile, D., Kazakou, E., Fortunel, C., Hummel, I., & Garnier, E. (2007). Let the concept of trait be functional! Oikos, 116(5), 882–892.

Weibull, A.-C., & Östman, Ö. (2003). Species composition in agroecosystems: The effect of landscape, habitat, and farm management. Basic and Applied Ecology, 4(4), 349–361.

Weibull, A.-C., Östman, Ö., & Granqvist, Å. (2003). Species richness in agroecosystems: The effect of landscape, habitat and farm management. Biodiversity & Conservation, 12(7), 1335–1355.

Winqvist, C., Bengtsson, J., Öckinger, E., Aavik, T., Berendse, F., Clement, L. W., Fischer, C., Flohre, A., Geiger, F., & Liira, J. (2014). Species’ traits influence ground beetle responses to farm and landscape level agricultural intensification in Europe. Journal of Insect Conservation, 18(5), 837–846.

Woodcock, B. A., Harrower, C., Redhead, J., Edwards, M., Vanbergen, A. J., Heard, M. S., Roy, D. B., & Pywell, R. F. (2014). National patterns of functional diversity and redundancy in predatory ground beetles and bees associated with key UK arable crops. Journal of Applied Ecology, 51(1), 142–151.

Zaller, J. G., Moser, D., Drapela, T., & Frank, T. (2009). Ground-dwelling predators can affect within-field pest insect emergence in winter oilseed rape fields. BioControl, 54(2), 247.

