## Appendix for "Functional traits of carabid beetles reveal seasonal variation in community assembly in annual crops"

**Appendix S1.**

**Table S1.** Description of the dataset.

| Year | Preserv. solution | Sampling date range (JJ) | NSes (mean) | Number of sampled fields/communities per crop | | | | |
| --- | --- | --- | --- | --- | --- | --- | --- | --- |
|  |  |  |  | Alfalfa | Grassland | OSR | Sun | WC |
| 2005 | EG | 91-225 | 1-7 (4.83) | 4/26 | 1/2 | 2/9 | 3/15 | 2/6 |
| 2006 | EG | 89-209 | 2-5 (3.93) | 3/10 | 3/13 | 4/16 | 1/3 | 3/13 |
| 2007 | EG | 147-174 | 1-2 (1.71) | 6/10 | 0/0 | 4/7 | 1/2 | 3/5 |
| 2008 | MPG | 150-178 | 2 | 5/10 | 4/8 | 4/8 | 2/4 | 4/8 |
| 2009 | EG/MPG | 114-212 | 1-5 (1.4) | 45/73 | 31/39 | 6/24 | 6/19 | 138/161 |
| 2010 | EG/MPG | 155-188 | 1-3 (1.09) | 52/58 | 31/33 | 4/7 | 3/4 | 104/109 |
| 2011 | Alcohol | 118-178 | 1-4 (2.34) | 14/32 | 4/8 | 13/29 | 12/32 | 13/30 |
| 2012 | Alcohol | 90-184 | 1-6 (4.04) | 12/43 | 4/15 | 14/46 | 8/28 | 39/179 |
| 2013 | Alcohol | 124-184 | 1-3 (1.51) | 7/13 | 0/0 | 9/14 | 3/4 | 24/34 |

Year: trapping year; Preservative solution: EG, ethylene-glycol, MPG, mono-propylene-glycol, Alcohol, ethanol; Sampling date range as Julian dates; NSes: number of trapping sessions per field (with mean value into brackets); Number of sampled fields per crop: perennial crops (Alfalfa, Grassland) and annual crops (OSR: oilseed rape; Sun: sunflower; WC: winter cereal).

**Appendix S2.** Species level traits

**Table S2.** Correlation between traits and PCA axis (r). Bold coefficients are significantly associated with PCA axis.

| Traits | PCA Axis 1 | PCA Axis 2 | PCA Axis 3 |
| --- | --- | --- | --- |
| *Wg* | 0.37 | **0.85** | 0.16 |
| *Bs* | **0.93** | 0.32 | -0.18 |
| *Lg* | **0.95** | 0.25 | 0.05 |
| *Fm* | **0.95** | 0.12 | -0.23 |
| *Wg:Bs* | -0.30 | **0.74** | 0.43 |
| *Fm:Tb* | **0.51** | 0.12 | 0.05 |
| *Fm:Bs* | -0.25 | **-0.67** | 0.19 |
| *Md:Hd* | **-0.87** | -0.36 | 0.10 |
| *MdLb* | 0.04 | 0.08 | **0.59** |
| *Bl:Bw* | **0.65** | -0.26 | 0.00 |
| *Hl:Hw* | -0.34 | **-0.56** | **0.56** |

**Table S3.** Correlation *r* among individual traits at the species level (see Fig. 2 and the materials and methods section for definitions of trait abbreviations). Positive (blue) and negative (red) correlations with *r* > 0.70 are in bold characters.

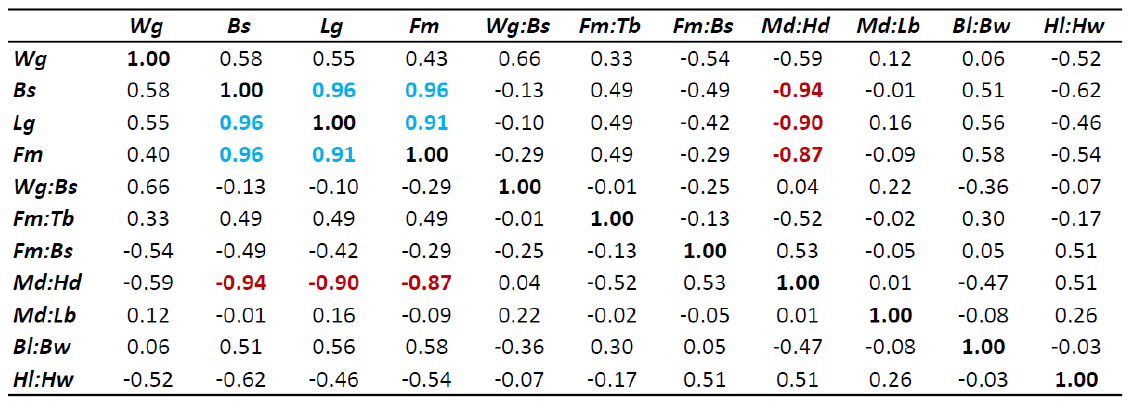

**Appendix S3.** Model selection

**Table S4.** Model selection table, indicating the relative accuracy of full models (grey lines) and their nine more statistically accurate and biologically meaningful sub-models. Models with ΔAIC < 4 are indicated in bold characters.

| **Response variable** | **Model _ID** | **Intercept** | **Year** | **Solution** | **Crop** | **JD** | **JD²** | **JD^3^** | **Crop:JD** | **Crop:JD²** | **Crop:JD^3^** | **df** | **logLik** | **AICc** | **delta** |
| --- | --- | --- | --- | --- | --- | --- | --- | --- | --- | --- | --- | --- | --- | --- | --- |
| CWM1 | **64** | **-107.9** | **0.054** | **+** | **+** | **-0.042** | **0.080** | **0.029** | **+** | **+** |  | **21** | **-427.5** | **898** | **0** |
|  | **128** | **-106.9** | **0.053** | **+** | **+** | **-0.043** | **0.080** | **0.029** | **+** | **+** | **+** | **25** | **-424.9** | **901.1** | **3.11** |
|  | 112 | -114.9 | 0.057 | + | + | -0.035 | 0.081 | 0.027 |  | + | + | 21 | -436.3 | 915.5 | 17.47 |
|  | 110 | -106.5 | 0.053 | + | + |  | 0.082 | 0.018 |  | + | + | 20 | -437.6 | 916 | 18.06 |
|  | 56 | -85.96 | 0.043 | + | + | 0.046 | 0.083 |  | + | + |  | 20 | -437.8 | 916.4 | 18.42 |
|  | 32 | -109.1 | 0.054 | + | + | -0.051 | 0.028 | 0.030 | + |  |  | 17 | -443.1 | 920.8 | 22.82 |
|  | 96 | -108.6 | 0.054 | + | + | -0.058 | 0.029 | 0.033 | + |  | + | 21 | -439.2 | 921.3 | 23.35 |
|  | 28 | -95.13 |  | + | + | -0.048 | 0.029 | 0.047 | + |  |  | 16 | -448.1 | 928.8 | 30.82 |
|  | 92 | -93.99 | 0.047 | + | + | -0.059 |  | 0.033 | + |  | + | 20 | -444.4 | 929.6 | 31.59 |
|  | 48 | -112.9 | 0.056 | + | + | -0.035 | 0.081 | 0.029 |  | + |  | 17 | -448.9 | 932.5 | 34.47 |
| CWM2 | **48** | **-32.6** | **0.016** | **+** | **+** | **-0.048** | **0.059** | **0.012** |  | **+** |  | **17** | **-329.8** | **694.3** | **0** |
|  | **40** | **-23.87** | **0.012** | **+** | **+** | **-0.013** | **0.060** |  |  | **+** |  | **16** | **-332.0** | **696.5** | **2.19** |
|  | 64 | -28.17 | 0.014 | + | + | -0.029 | 0.060 | 0.011 | + | + |  | 21 | -327.7 | 698.3 | 4.04 |
|  | 56 | -19.64 |  | + | + | 0.005 | 0.061 | 0.010 | + | + |  | 20 | -329.4 | 699.7 | 5.43 |
|  | 128 | -28.75 | 0.014 | + | + | -0.017 | 0.061 | 0.007 | + | + | + | 25 | -324.6 | 700.5 | 6.27 |
|  | 112 | -31.56 | 0.016 | + | + | -0.048 | 0.059 | 0.014 |  | + | + | 21 | -329.5 | 701.8 | 7.52 |
|  | 16 | -31.56 | 0.016 | + | + | -0.049 | 0.066 | 0.013 |  |  |  | 13 | -343.6 | 713.6 | 19.27 |
|  | 32 | -26.75 | 0.013 | + | + | -0.030 | 0.063 | 0.012 | + |  |  | 17 | -340.3 | 715.2 | 20.97 |
|  | 8 | -22.83 | 0.011 | + | + | -0.013 | 0.066 |  |  |  |  | 12 | -345.8 | 715.9 | 21.62 |
|  | 24 | -18.27 | 0.009 | + | + | 0.005 | 0.063 |  | + |  |  | 16 | -342.2 | 716.9 | 22.6 |
| CWM3 | **128** | **7.645** | **-0.004** | **+** | **+** | **-0.029** | **-0.028** | **-0.011** | **+** | **+** | **+** | **25** | **-583.1** | **1217.6** | **0** |
|  | 96 | 6.575 | -0.003 | + | + | -0.029 | -0.022 | -0.011 | + |  | + | 21 | -593.4 | 1229.6 | 12.06 |
|  | 64 | 13.62 | -0.007 | + | + | -0.093 | -0.032 | 0.011 | + | + |  | 21 | -593.6 | 1230 | 12.44 |
|  | 56 | 22.36 | -0.011 | + | + | -0.059 | -0.031 |  | + | + |  | 20 | -594.8 | 1230.3 | 12.76 |
|  | 32 | 12.21 | -0.006 | + | + | -0.095 | -0.026 | 0.012 | + |  |  | 17 | -605.5 | 1245.6 | 28.04 |
|  | 24 | 21.04 | -0.010 | + | + | -0.059 | -0.027 |  | + |  |  | 16 | -606.8 | 1246.2 | 28.61 |
|  | 20 | 8.536 | -0.004 | + | + | -0.060 |  |  | + |  |  | 15 | -610.3 | 1251.1 | 33.52 |
|  | 112 | 10.69 | -0.005 | + | + | -0.105 | -0.033 | 0.008 |  | + | + | 21 | -607.7 | 1258.3 | 40.7 |
|  | 48 | 8.599 | -0.004 | + | + | -0.100 | -0.033 | 0.011 |  | + |  | 17 | -617.0 | 1268.6 | 51.04 |
|  | 40 | 16.71 | -0.008 | + | + | -0.068 | -0.032 |  |  | + |  | 16 | -618.2 | 1268.8 | 51.28 |
| CWV1 | **112** | **83.09** | **-0.041** | **+** | **+** | **0.039** | **-0.078** | **-0.013** |  | **+** | **+** | **21** | **-26.4** | **95.6** | **0** |
|  | **128** | **83.6** | **-0.041** | **+** | **+** | **0.030** | **-0.079** | **-0.011** | **+** | **+** | **+** | **25** | **-23.0** | **97.4** | **1.72** |
|  | 64 | 78.91 | -0.039 | + | + | 0.031 | -0.077 | -0.012 | + | + |  | 21 | -29.7 | 102.3 | 6.67 |
|  | 80 | 78.31 | -0.039 | + | + | 0.039 | -0.043 | -0.013 |  |  | + | 17 | -34.0 | 102.6 | 6.98 |
|  | 96 | 80.48 | -0.040 | + | + | 0.041 | -0.042 | -0.013 | + |  | + | 21 | -30.4 | 103.8 | 8.13 |
|  | 56 | 69.94 | -0.035 | + | + | -0.005 | -0.078 |  | + | + |  | 20 | -33.0 | 106.9 | 11.23 |
|  | 48 | 84.61 | -0.042 | + | + | 0.040 | -0.078 | -0.014 |  | + |  | 17 | -36.3 | 107.2 | 11.54 |
|  | 32 | 74.73 | -0.037 | + | + | 0.036 | -0.040 | -0.012 | + |  |  | 17 | -37.7 | 109.9 | 14.27 |
|  | 40 | 75.25 | -0.037 | + | + | 0.002 | -0.079 |  |  | + |  | 16 | -40.4 | 113.3 | 17.65 |
|  | 24 | 65.99 | -0.033 | + | + | 0.000 | -0.039 |  | + |  |  | 16 | -41.1 | 114.6 | 19.01 |
| CWV2 | **128** | **65.02** | **-0.032** | **+** | **+** | **0.089** | **-0.055** | **-0.015** | **+** | **+** | **+** | **25** | **-117.5** | **286.3** | **0** |
|  | 48 | 66.17 | -0.033 | + | + | 0.119 | -0.053 | -0.022 |  | + |  | 17 | -128.0 | 290.6 | 4.36 |
|  | 64 | 63.32 | -0.031 | + | + | 0.109 | -0.053 | -0.022 | + | + |  | 21 | -124.0 | 290.9 | 4.67 |
|  | 96 | 62.63 | -0.031 | + | + | 0.093 | -0.038 | -0.016 | + |  | + | 21 | -126.7 | 296.3 | 10.04 |
|  | 112 | 66.09 | -0.033 | + | + | 0.120 | -0.053 | -0.023 |  | + | + | 21 | -127.6 | 298.1 | 11.86 |
|  | 32 | 60.45 | -0.030 | + | + | 0.111 | -0.036 | -0.022 | + |  |  | 17 | -133.9 | 302.3 | 16.08 |
|  | 16 | 63.1 | -0.031 | + | + | 0.120 | -0.038 | -0.022 |  |  |  | 13 | -138.6 | 303.6 | 17.38 |
|  | 40 | 51.04 | -0.025 | + | + | 0.056 | -0.055 |  |  | + |  | 16 | -137.7 | 307.9 | 21.65 |
|  | 56 | 47.1 | -0.023 | + | + | 0.042 | -0.055 |  | + | + |  | 20 | -133.6 | 308 | 21.79 |
|  | 80 | 62.59 | -0.031 | + | + | 0.120 | -0.039 | -0.023 |  |  | + | 17 | -138.1 | 310.8 | 24.58 |
| CWV3 | **96** | **-18.1** | **0.009** | **+** | **+** | **0.059** | **-0.010** | **-0.021** | **+** |  | **+** | **21** | **-212.9** | **468.6** | **0** |
|  | **128** | **-17.46** | **0.009** | **+** | **+** | **0.052** | **-0.029** | **-0.020** | **+** | **+** | **+** | **25** | **-209.6** | **470.4** | **1.81** |
|  | 80 | -18.03 | 0.009 | + | + | 0.045 | -0.010 | -0.018 |  |  | + | 17 | -219.4 | 473.5 | 4.87 |
|  | 16 | -18.52 | 0.009 | + | + | 0.046 | -0.007 | -0.016 |  |  |  | 13 | -224.0 | 474.3 | 5.7 |
|  | 48 | -16.78 | 0.009 | + | + | 0.047 | -0.029 | -0.016 |  | + |  | 17 | -220.1 | 474.8 | 6.15 |
|  | 15 | -34.19 | 0.017 | + |  | 0.035 | -0.010 | -0.013 |  |  |  | 9 | -228.5 | 475.2 | 6.56 |
|  | 112 | -15.99 | 0.008 | + | + | 0.046 | -0.030 | -0.018 |  | + | + | 21 | -216.3 | 475.5 | 6.85 |
|  | 1 | -46.51 | 0.023 | + |  |  |  |  |  |  |  | 6 | -232.4 | 476.9 | 8.23 |
|  | 2 | -33.43 | 0.017 | + | + |  |  |  |  |  |  | 10 | -228.7 | 477.7 | 9.06 |
|  | 3 | -46.86 | 0.023 | + |  | -0.002 |  |  |  |  |  | 7 | -232.4 | 478.8 | 10.22 |

**Appendix S4.** Relationship between carabid beetle morphological traits and feeding niche

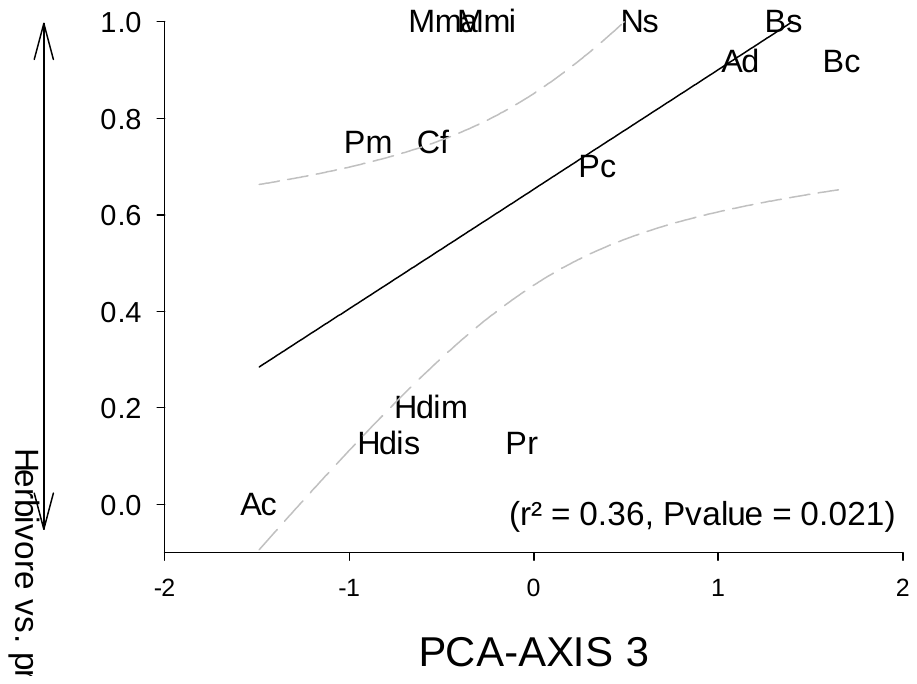

**Fig. S1.** Relationship between species traits (PCA axis 3) (see Appendix S2 and Fig. 2) and feeding niche of carabid species.

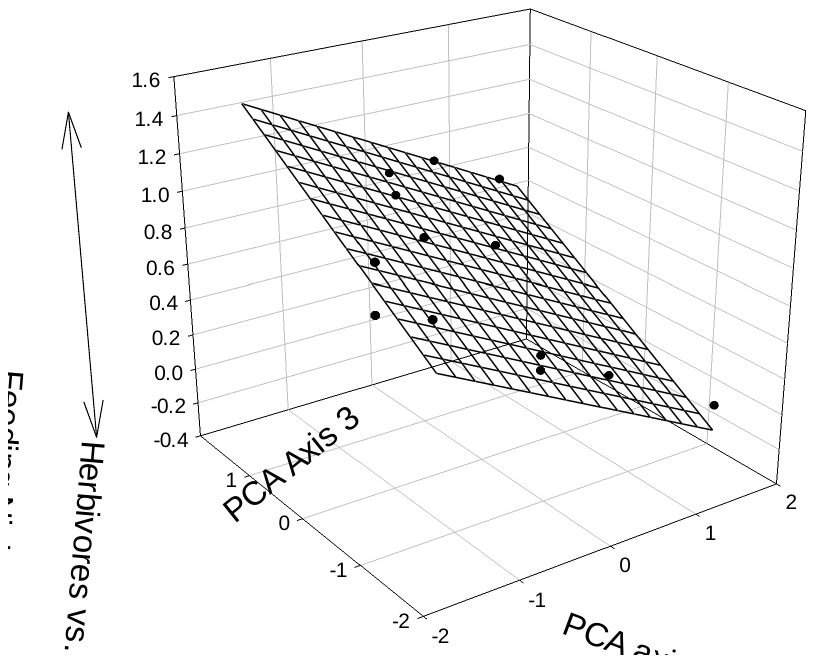

**Fig S2.** Relationship between PCA axes 2 and 3 (see Appendix S2 and Fig. 2) and feeding niche of carabid species. Each black dot represents one of the 13 species.
